# Excessive deliberation in social anxiety

**DOI:** 10.1101/522433

**Authors:** Lindsay E. Hunter, Elana A. Meer, Claire M. Gillan, Ming Hsu, Nathaniel D. Daw

## Abstract

A goal of computational psychiatry is to ground symptoms in more fundamental computational mechanisms. Theory suggests that rumination and other symptoms in mood disorders reflect dysregulated mental simulation, a process that normally serves to evaluate candidate actions. If so, these covert symptoms should have observable consequences: excessively deliberative choices, specifically about options related to the content of rumination. In two large general population samples, we examined how symptoms of social anxiety disorder (SAD) predict choices in a socially framed reinforcement learning task, the Patent Race game. Using a computational learning model to assess learning strategy, we found that self-reported social anxiety was indeed associated with an increase in deliberative evaluation. The effect was specific to learning from a particular (“upward counterfactual”) subset of feedback, broadly matching the biased content of rumination in SAD. It was also robust to controlling for other psychiatric symptoms. These results ground the symptoms of SAD, such as overthinking and paralysis in social interactions, in well characterized neuro-computational mechanisms and offer a rare example of enhanced function in disease

## Introduction

Unique among areas of medicine, psychiatry has no laboratory diagnostic tests. This is largely due to a lack of understanding how mental health symptoms arise from dysfunction in underlying brain mechanisms. Recent research has attempted to fill this gap by connecting these disorders to relatively well characterized neuro-computational systems, notably those that support reinforcement learning (RL) (Maia & Frank, 2011; Huys et al., 2012, 2016; Montague et al., 2012; Moutoussis et al., 2017). In this respect, a key feature of RL in the brain is that it arises from a combination of at least two evaluative mechanisms, more deliberative vs. automatic, which have been formalized in terms of model-based and model-free learning (Daw et al., 2005). Model-based learning evaluates actions by iteratively simulating their consequences using a learned representation, or “model” of the task’s contingencies, such as a spatial map. Model-free learning skirts this computation by learning actions’ long run values directly from experience when they are chosen; these values permit quick but inflexible decisions and are a potential substrate for habits. A line of research describes the biological substrates for these functions, such as representations of future spatial trajectories in hippocampus that may support mental simulations of candidate routes for model-based evaluation (Johnson & Redish, 2007; Mattar & Daw, 2018) and dopaminergic temporal-difference prediction error signals suited for model-free learning (Schultz et al., 1997). Although adaptive behavior relies on the ability to flexibly recruit both strategies, people also vary greatly in their tendency to do so. Accordingly, by comparing RL models to trial-by-trial choices in people learning sequential choice tasks (e.g., two-step Markov decision processes, MDPs; Daw et al., 2011), the degree to which subjects utilize model-based RL has been shown to vary both situationally (e.g., under dual-task interference; Otto et al., 2013) and between individuals (e.g., with genotypic variations that affect prefrontal dopamine; Doll et al., 2016).

Abnormal imbalance between these mechanisms has also been the focus of persistent speculation regarding mental illness. In particular, it has long been suggested that symptoms related to compulsion (a dimension that cuts across illnesses including OCD and drug abuse) might arise from an imbalance that favors automaticity (Everitt & Robbins, 2005; Gillan et al., 2011, 2014, 2016; Reiter et al., 2017). More tentatively, theorists have suggested that some aspects of mood disorders (notably, rumination in depression or excessive caution and overthinking in anxiety) might relate to a converse imbalance, favoring *excess* deliberation (Huys et al., 2012; Huys et al., 2015; Solway et al., submitted). Indeed, for a variety of disorders involving compulsion, both patients’ diagnoses (Voon et al., 2015), and self-reported symptoms in a large general-population sample (Gillan et al., 2016), are associated with deficient model-based learning in a two-step MDP. There is as yet less evidence relating depression or anxiety to increased simulation of action-outcome contingencies in value-based learning, though some analyses in the Gillan et al. (2016) dataset revealed a small trend of increased model-based evaluation specifically for social anxiety, rather than other mood disorders. Social anxiety is also an interesting test case, both practically (because clinically significant levels of it are frequent in the population tested online; Shapiro et al., 2013) and substantively, because it involves enhanced mentalizing and counterfactual thinking (see Norton & Abbott, 2016 for a review), both psychological constructs closely related to model-based evaluation.

We thus sought to investigate the hypothesis that social anxiety is associated with increased deliberation. To better probe their relationship, we turned to another well-studied type of learning task with a more social framing: a competitive economic game (Zhu et al., 2012; Rapoport & Amaldoss, 2000), the patent race. As with MDPs, choices on this sort of task are well-characterized by a subject-specific combination of two strategies, paralleling model-free and model-based RL: direct (model-free) learning about which moves are successful, vs. deriving moves’ values indirectly based on which moves the opponent prefers (equivalent to a model of the opponent; Camerer & Ho, 2000). Moreover, the patterns of neuroimaging correlates and individual differences (e.g., with dopaminergic genes) related to this dichotomy parallel those reported for MDPs (Zhu et al., 2012; Set et al., 2014).

The patent race game also enables us to investigate anxiety’s effects not just on overall model usage, but also on more granular operations of planning. Recent research aims to decompose model-based evaluation into a series of steps, such as simulating individual candidate actions or accessing memories for particular events (Cushman & Morris, 2015; Mattar & Daw, 2018). Meanwhile, rumination is characterized by a narrow preoccupation with particular thoughts. If indeed it constitutes runaway model-based evaluation, these content biases should manifest in a specific tendency to update values for certain actions rather than others.

In social anxiety, rumination includes an excess of “upward counterfactual” thoughts about previous social interactions: that is, “if only” thoughts about how the events could have gone better, which fuels anticipatory anxiety of future social interactions (Kocovski et al., 2005, 2015). Model-based evaluation in the patent-race task can also be expressed in terms of counterfactual updating – computing the value of moves not taken, given the opponent’s moves (Camerer and Ho, 1999) – and a bias toward *upward* counterfactuals would predict a tendency toward model-based updating for a subset of the six moves available. If such a bias is observed in choices, this would speak both to the granular mechanism of planning, and the effects of rumination on it.

Accordingly, to examine the relationship between social anxiety and model-based deliberation, we conducted two large-scale, online experiments. In each experiment, 500 subjects completed the Liebowitz Social Anxiety Scale (LSAS) and played 80 rounds of a patent race game against a computerized opponent. In the second cohort, we also assessed symptoms for a broader range of psychiatric symptoms, allowing us to probe the specificity of our findings.

## Methods

### Participants & Procedures

Overall, 1000 participants (500 per experiment) were recruited online using Amazon’s Mechanical Turk (AMT), of whom 966 (N_1_ = 489; N_2_ = 477) provided a complete dataset. Participants were paid a base rate in addition to a bonus proportional to their (nominal) earnings during the reinforcement-learning task. Subjects were based in the USA (i.e. had a US billing address with an associated US credit card, debit card or bank account), 95% of their previous tasks were approved, and were 18 years or older. All procedures were pre-approved by Princeton University’s Institutional Review Board.

Via their web browser, all participants completed the Liebowitz Social Anxiety Scale (LSAS), Liebowitz, 1987), an abbreviated nine-item Ravens’ matrix test to approximate IQ (Bilker et al., 2012), and 80 rounds of a patent race game (Zhu et al., 2012). Procedures for the two experiments differed only in that subjects in Experiment 2 completed a more comprehensive psychopathological assessment (in addition to LSAS) before proceeding to the IQ test and patent race game. In particular, Experiment 2 included a battery of 209 multiple choice questions which gauged symptom severity across a range of disorders and constructs (the same battery used by Gillan et al., 2016; Renault et al, 2018): alcoholism (Alcohol Use Disorders Identification Test), apathy (Apathy Evaluation Scale), impulsivity (Barratt Impulsiveness Scale 11), eating disorders (Eating Attitudes Test), social anxiety (Liebowitz Social Anxiety Scale), obsessive-compulsive disorder (OCD) (Obsessive-Compulsive Inventory-Revised [OCI-R]), schizotypy (Short Scales for Measuring Schizotypy), depression (Self-Rating Depression Scale), and generalized anxiety (State Trait Anxiety Inventory).

### Patent Race Game

Subjects played 80 rounds of an asymmetric patent race task, a competitive, simultaneous move game in which a ‘strong’ player with more resources competes with a ‘weak’ player with fewer resources (Rapoport and Amaldoss, 2000; Zhu et al., 2012). In each round of the game, subjects (who played the “weak” role) were endowed with $4 and chose how much to invest (in integer dollars, $0 - $4) to obtain a $10 prize. The computerized opponent was endowed $5 and thus held a stronger position. The rules of the game, including each player’s endowments and the payoffs conditional on different moves, were common knowledge and remained fixed throughout the game. When a player invested strictly more than their opponent, they won the $10 prize on that round. (In case of tie, no one received the prize.) Regardless of the outcome, players kept the uninvested portion of their endowment.

Subjects were advised that the computerized opponent’s choices on each round were drawn from a pool of choices made by previous human participants at that round of the game. Thus, although opponents were anonymous and unlikely to be encountered more than once in a row, they represented people at the same stage of progression through the task as the subject.

### Learning models

Since the distribution of opponents’ moves was unknown to the subjects (and potentially nonstationary), the game presents a learning problem: finding which moves are most effective. Two leading models for this process in behavioral game theory correspond to model-free and model-based RL (though model-free RL is known in this literature simply as “reinforcement learning” while model-based RL is referred to as “belief learning”); a third model, known as Experience-Weighted Attraction (EWA; Camerer & Ho, 1999) characterizes behavior by a weighted combination of these two strategies.

The model-free rule is simple Q-learning – it maintains an expected value for each possible move, updated whenever a move is chosen according to the received payoff (Erev & Roth 1998). In its original formulation (Brown, 1951; Cheung & Friedman, 1997), belief learning turns on learning the opponent’s move distribution (a model about the opponent’s preferences or “beliefs”, updated each time their move is observed). With this and the payoff matrix, the expected payoffs for each of the players’ responses can be computed. In fact, marginalizing the beliefs, the same payoff estimates for the player’s moves can be updated in place at each timestep, by updating each of them according to the reward that *would have* been received had the player chosen that move, given the opponent’s move (Camerer & Ho, 1999). This approach can be viewed either as an algebraic trick for conveniently implementing the predictions of the model-based rule; or as a substantive hypothesis for how these computations might actually be implemented in the brain using counterfactual updates in place of the belief model. Similar approaches, which substitute replayed experience for a world model, have also been examined for other RL tasks like spatial navigation (Sutton, 1991; Mattar & Daw, 2018)

In the context of this game, model-free and model-based learning make different predictions about how the subjective value of each strategy (i.e., each possible investment amount) is updated with experience at each round. Consider a round in which the subject invests $4 and the opponent invests $2. Here the subject’s choice to invest $4 results in a total return of $10. Model-free learning would update the expected value of investing $4 by moving it closer to $10. In belief learning (implemented via counterfactual updating) the subject further updates the value of each other move by calculating the return it would have yielded on the previous trial given the observed investment made by the opponent. For example, in this case, the subject will update the value of investing $2 toward $2 and the value of $3 toward $11, since these are the amounts is the amount the subject would have won given that the opponent invested $2.

### Computational Modeling

Following previous work (Rapoport and Amaldoss 2000; Amaldoss and Jain, 2002; Zhu et al., 2012), we modeled subjects’ learning on this task using Camerer and Ho’s (1999) EWA learning model. This constitutes a weighted combination of the two learning strategies discussed above, analogous to hybrid model-based/free models previously used for human choices in MDPs (Daw et al., 2011). Because of the equivalence between model updating and counterfactual updating in this class of tasks, the weighted combination of both strategies can be expressed in a single update rule comprising weighted updates from experienced and counterfactual rewards.

In particular, at round *t*, a player updates their estimate of the value *V*, for each move *k* according to:

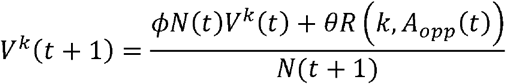

where

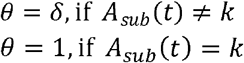

with free parameters *ϕ* and *δ*. In these expressions, *A*_*sub*_ and *A*_*opp*_ denote the moves chosen by the subject and the opponent (i.e., $0 - $4 and $0 - $5), and *R* is the payoff matrix giving the subject’s reward as a function of both moves. At each step, the subject also updates an experience counter *N*, with free parameter *ρ*:

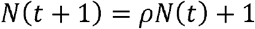

Choices were taken as softmax in *V*^*k*^(*t*), i.e.

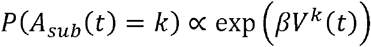

with free inverse temperature parameter *β*.

We take *N*(1) = 1 and introduce free parameters for the initial values *V*^*k*^(1) (for *k* = 0 - 4), capturing any *a priori* preferences for the actions.

Although the notation is unusual, readers familiar with standard RL models will recognize that for *ρ* = 0 (hence, *N*(*t*) = 1 for all *t*), *δ* = 0, and a learning rate of (1 - *ϕ*) the value update reduces to a standard delta rule in the chosen option and received reward, but with the *R* (and hence all *V*^*k*^) rescaled by 1/(1 − *ϕ*). This scaling can be canceled by inversely scaling *β* by the same factor, making the model equivalent to a standard model-free Q-learning update. For *ρ* > 0, *N* accumulates and drives a sort of learning rate decay. For simplicity, and following previous work (Set et al., 2014) we took the two decay parameters as equal, i.e. *ρ* = *ϕ*.

Our main interest is the parameter *δ* in the original model, which controls the relative weight of counterfactual updating (*δ* = 0, fully model-free; *δ* = 1, fully model-based), and *δ*^+^ in the variant model, which isolates the part of the effect due to upwards counterfactual updating. In total, the model contained 8 free parameters.

To account for the distinction between “upward” and “downward” counterfactual, we considered an additional variant of the EWA model in which, *δ*, the degree of counterfactual updating, is valence-dependent. This parameterization splits the free parameter *δ* into *δ*^+^ and *δ*^−^, which control upwards- and downwards- counterfactual learning, respectively. The reward received on a given round, *R*(*A*_*sub*_(*t*),*A*_*opp*_(*t*)), serves as the reference point. Actions (investments) that would have resulted in more reward than what was won in reality (based on opponents’ investment on that round), are updated in proportion to 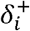 whereas actions that would have resulted in less reward are weighted instead by 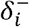.

Accordingly, the update rule for values is the same as before, but with

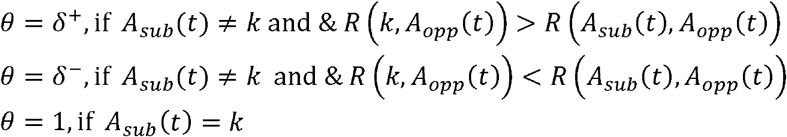

Note that we recover the original model when *δ*^+^ = *δ*^−^. In total, the valence-dependent model contains 9 free parameters.

We estimated the free parameters of each model per subject using an EM optimization algorithm (Huys et al., 2011) implemented in the Julia language (Bezanson et al., 2012). This procedure maximizes the joint likelihood of each individual’s sequence of choices, where each individual’s parameter estimates are random effects drawn from group-level Gaussian parameter distributions, whose means and variances are also estimated. We first used values derived separately for each experiment in to ensure that our major patterns of results held within Experiments 1 and 2 independently (*SI Figure 7, SI Tables 3 & 4*), and then pooled the data across Experiments 1 and 2.

We used a set of multiple linear regressions, one for each parameter, to test whether LSAS scores or symptom factor scores (see below) predicted each of the free parameters from the EWA models. These models included an additional covariate to control for Ravens’ matrices scores, using the score predicted for the full 60-item set based on the 9 problems given (Bilker et al., 2012). All predictors were standardized (z-scored) for interpretability of coefficients. Auxiliary analyses also considered age as a predictor, but this had no appreciable effects on the results. To test whether the effects were different between parameters *δ*^+^ and *δ*^−^, we repeated the regression using a linear mixed-effects model to capture the repeated-measure structure.

To assess the relative fit of the elaborated vs. standard variant of the EWA model while correcting for overfitting due to both group- and subject-level parameters, we computed integrated Bayesian information criterion (iBIC) scores (Huys et al., 2012). This was defined as the marginal likelihood of the data given either model, aggregated across subjects, marginalizing per-subject parameters with the Laplace approximation, and penalizing for the group-level parameters using BIC (Huys et all, 2012).

### Factor Analysis

For Experiment 2, we used the factors identified by Gillan et al. (2016) to reduce responses on the 209 items from the nine psychiatric symptom scales to scores on three dimensions that capture much of the intersubject variance. These were labeled by Gillan et al. (2016), as ‘Anxious-Depression’, ‘Compulsive Behavior and Intrusive Thought’ and ‘Social Withdrawal’ based on the items with the strongest loadings for each factor (*SI Figure 6*). We verified that the factor analysis procedure described by Gillan et al. (2016), when applied to our data, produced substantially the same factor structure (correlations between factor loadings: Factor 1: R = .94, p <1e-96; Factor 2: R = .91, p <1e-79; Factor 3: R = .91, p <1e-80). Because Gillan’s study analyzed the same battery of questionnaires using a much larger sample (N=1,413) we used the factor loadings estimated in that study to construct factor scores for each subject.

## Results

In line with recommendations for studies conducted using Amazon’s Mechanical Turk (AMT), a priori exclusion criteria were applied to ensure data quality (Crump et al., 2013). Of the 966 participants who completed the task, we eliminated participants who shirked either the Ravens matrix test (N_1_=36; N_2_= 49) or the patent race task (N_1_=41; N_2_=97) (77 & 146 were removed from experiment 1 and experiment 2 respectively, where the larger number in the second study is likely due to the longer session). Specifically, we removed from consideration subjects who got 0 or 1 items correct on Raven’s matrices or who chose the same move on more than 95% of trials (i.e. >76/80 rounds) in the patent race (Zhu et al., 2012). The remaining analyses concern the data of 743 participants (N_1_=412; N_2_=331).

Consistent with previous reports for the AMT population (Shapiro et al., 2013), LSAS scores were high (M=50; SD=30, out of 140), with the average participant (and 72% of all participants) above a standard threshold (30) for mild clinically significant effects (*Figure 2*). On average, subjects answered 5.4 (SD=1.8) of the nine Ravens’ matrix problems correctly, which corresponds to a predicted mean of 45 (SD=8.4) correct on the full 60-question set (using the weighted prediction from Bilker et al., 2012) (Figure 2). LSAS and IQs scores were distributed similarly across subjects in Experiments 1 and 2 (*SI Figures 2 & 3*).

**Figure 1:**
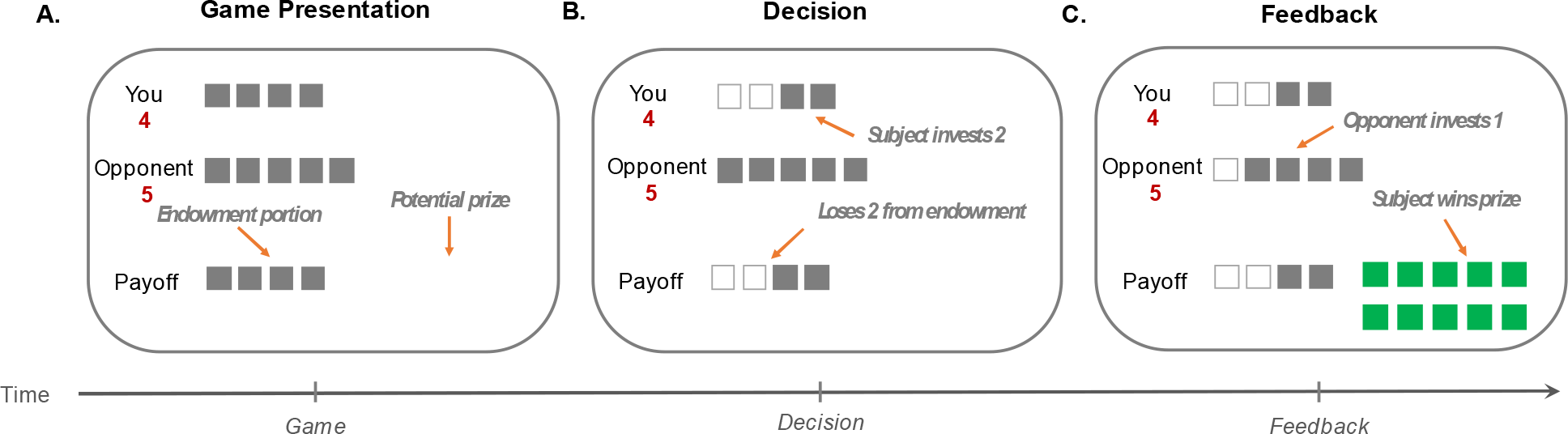
Schematic of Patent Race Game. Prior to each round, a fixation screen appeared for a random duration between 4–8 s. A.) Subjects were then presented with information regarding their endowment ($4), the endowment of the opponent ($5), and the potential prize ($10). B.) The arrow keys allowed subjects to select how much to invest (indicated by the number of white boxes) and the space bar was then used to submit the selected investment amount. C.) The opponent’s choice was revealed 2–6 s later. If the subject’s investment was strictly more than those of the opponent, the subject won the prize; In either case, the subject kept the portion of the endowment not invested.

**Figure 2:**
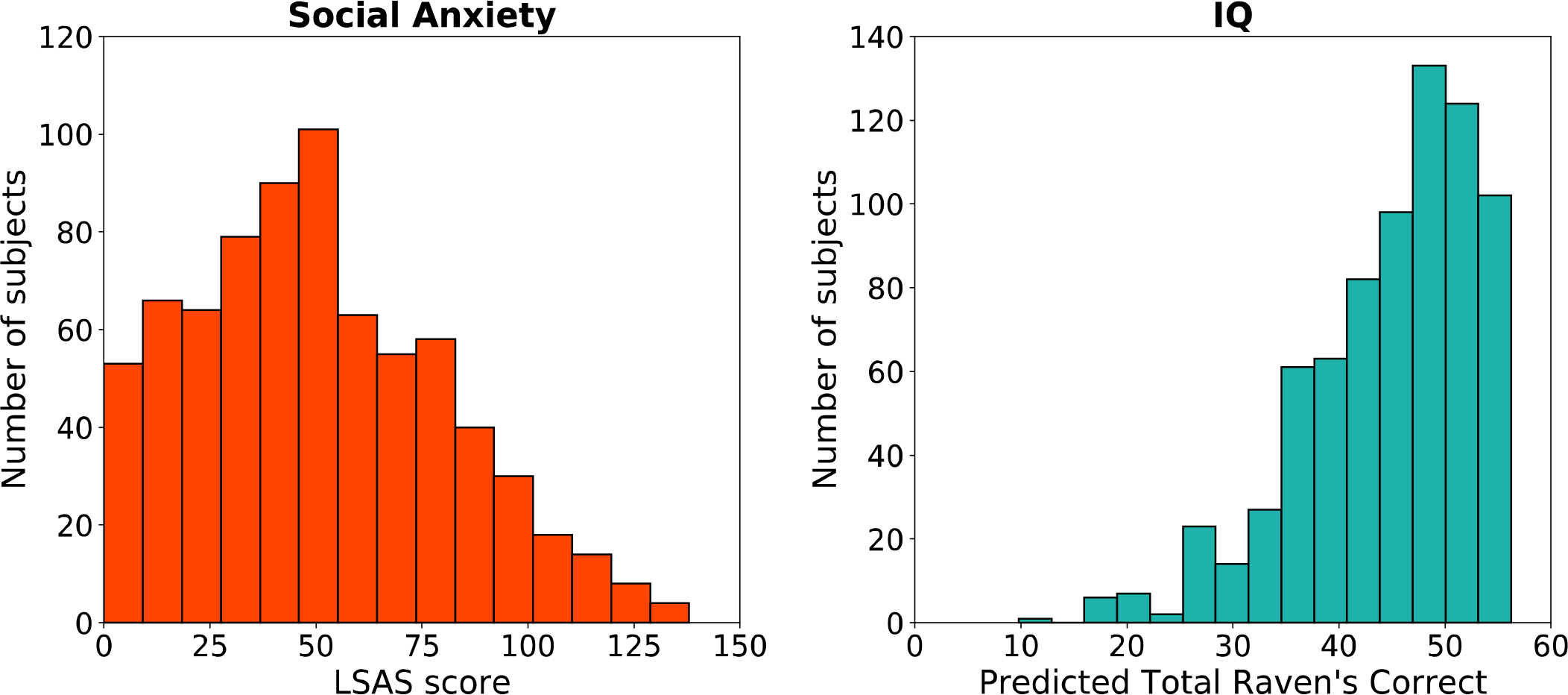
Distributions of self-report social anxiety and IQ scores for Experiments 1 and 2 (N= 743). Social Anxiety scores are based on Liebowitz Social Anxiety Scale (LSAS), (left) and IQ scores are based on subjects’ responses on the 9- item abbreviated version of the Raven’s Standard Progressive Matrices (RSPM) (right).

Behavior in the patent race game was consistent with patterns of investment behavior observed in previous studies featuring the same task as were estimated values for the EWA parameters (Zhu et al., 2012) (*SI Table 8*). Our main hypothesis was that higher self-reported LSAS scores would be associated with larger fit values of the parameter *δ*, corresponding to increased reliance on model-based, counterfactual updating. Indeed, the two variables were positively related (*SI Figure 7, SI Tables 3 & 4*) i.e., higher self-reported social anxiety predicted greater use of counterfactual updating in the patent race task in each experiment (t_412_ = 2.59, p = 0.01; t_331_ = 2.507, p = 0.01). We also found that the parameters did not differ between the two experiments, and that the positive influence of *δ* on social anxiety thus reached an even higher degree of significance (t_740_ = 3.63, p = 0.0003) when the data were pooled across experiments for visualization. (*Table 1*). For convenience we pool the datasets for the analyses below, but we report all analyses broken down by experiment in the supplemental material. The relationship between social anxiety and *δ* was selective; no significant correlations were observed between LSAS and the other parameters (*Figure 3*). These results also control for any effects of the Ravens matrix IQ score, which is included as an additional explanatory variable. As illustrated in Figure 3, intelligence was related to the model parameters, but with a very different pattern than LSAS. In particular, every 1 SD increase in IQ predicted a 7% increase in inverse temperature β (p<.001), consistent with the intuition that higher intelligence leads to improvement in generic task performance.

**Table 1:** Multivariate linear regressions of social anxiety (LSAS) and IQ (Ravens Matrices Scores) on estimated EWA parameters. Here *β* represents the inverse temperature, *ϕ* controls the learning rate, and *δ* dictates the rate of counterfactual updating.

| DV | IV | Est. (SE) | t-value | Pr>\|t\| |
| --- | --- | --- | --- | --- |
| $\beta$ | Intercept | 1.182 (0.020) | 59.8 | <0.001*** |
|  | Social Anxiety | -0.024 (0.020) | -1.21 | 0.228 |
|  | IQ | 0.084 (0.020) | 4.26 | <0.001*** |
| $\phi$ | Intercept | 0.836 (0.006) | 132 | <0.001*** |
|  | Social Anxiety | -0.003 (0.006) | -0.420 | 0.675 |
|  | IQ | -0.010 (0.006) | -1.61 | 0.107 |
| $\delta$ | Intercept | 0.217 (0.008) | 26.9 | <0.001*** |
|  | Social Anxiety | 0.029 (0.008) | 3.61 | <0.001*** |
|  | IQ | 0.011 (0.008) | 1.42 | 0.156 |

**Figure 3:**
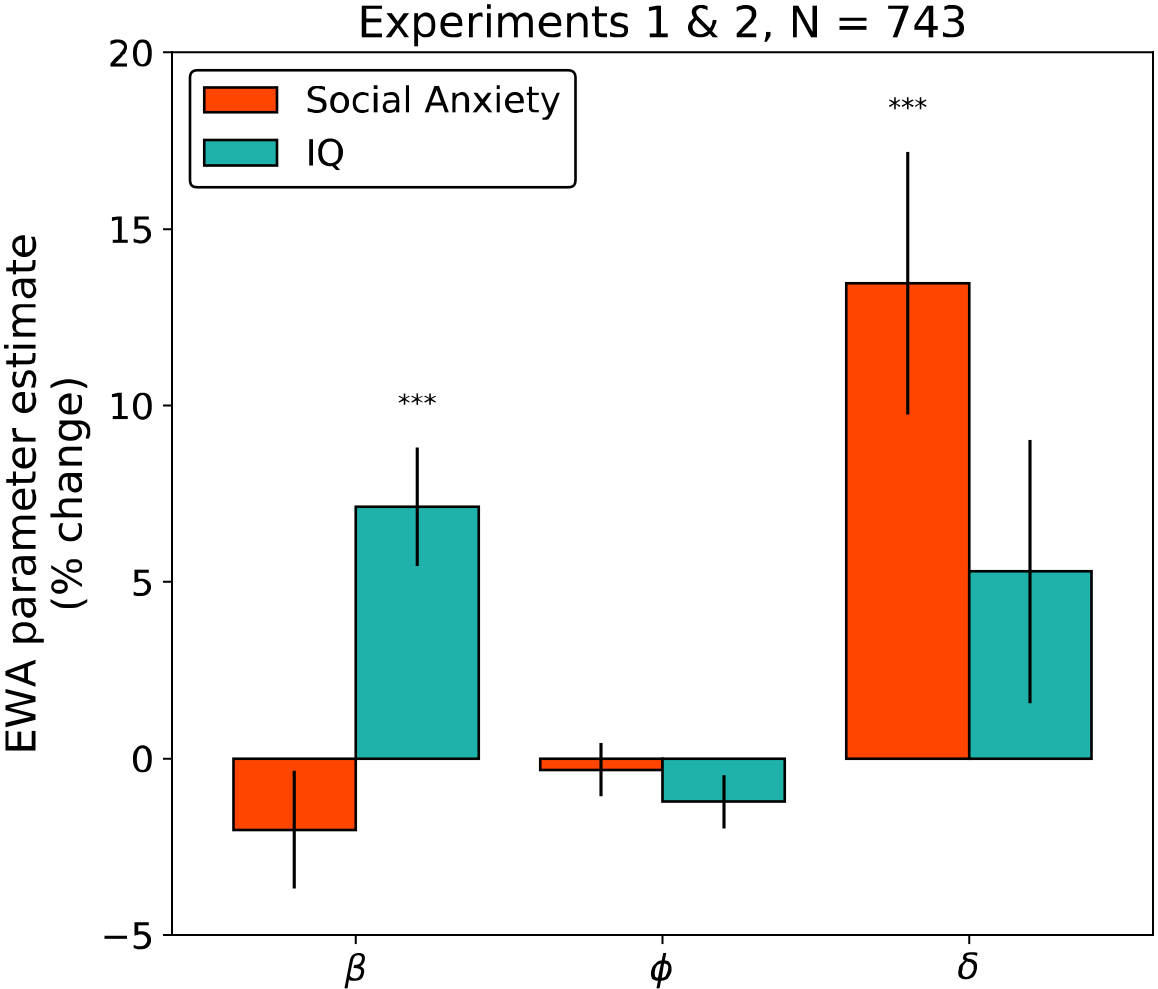
Percent change in EWA parameters as a function of social anxiety (LSAS) and IQ (Abbreviated 9-item Raven’s Matrices) for subjects from Experiments 1 and 2 (N=743). Here *β* represents the inverse temperature, *ϕ* controls the learning rate, and *δ* dictates the rate of counterfactual updating. The y-axes indicate the percent change in the parameter for each change of 1 standard deviation (SD) in the predictor. Error bars indicate 1 standard error.

Next, we used an elaborated variant of the EWA model to investigate the hypothesis that the effects of social anxiety would be specifically driven by counterfactual updating about subset of options. This model subdivides the parameter δ into two, *δ*^+^ and *δ*^−^, which govern counterfactual updating separately for options that would have been better, or respectively worse, than the one taken. We first verified that the elaborated model fit choices better than the original one, accounting for overfitting from the additional free parameters. Indeed, the iBIC score for the extended variant was lower (indicating a better fit) than that of the standard EWA model for the merged dataset (*iBIC*^*δ*^ = 57511, 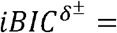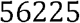) as well as for each individual experiment 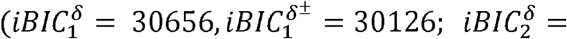 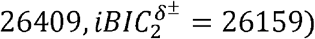.

If increased deliberation in social anxiety is driven by rumination about the events in the task, then the behavioral effects should reflect the biases of that rumination. In particular, post-event processing in people with high social anxiety involves a predominance of upward counterfactual thoughts (‘if only’ thoughts about how the situation could have gone *better*; Kocovski, Endler, Rector, & Flett, 2005; Brozovich et al., 2008). This suggests the link between social anxiety and *δ* is mediated by a more specific relationship between social anxiety and *δ*^+^ (upwards counterfactual updating). Indeed, social anxiety predicted a robust increase in upwards counterfactual updating, indexed by *δ*^+^ (p<0.0001), but had no significant relationship with *δ*^−^ (*Figure 4, Table 2*). Again, this result was significant in each experiment considered separately (*SI Figure 7, SI Tables 3 & 4*); and again, the pattern of correlations with IQ was different and less selective. We also verified that the association between social anxiety and *δ*^+^ was significantly greater than that for *δ*_−_ it was (p<.05 exp1, p<.022 exp2, *p* < .0014, combined data).

**Figure 4:**
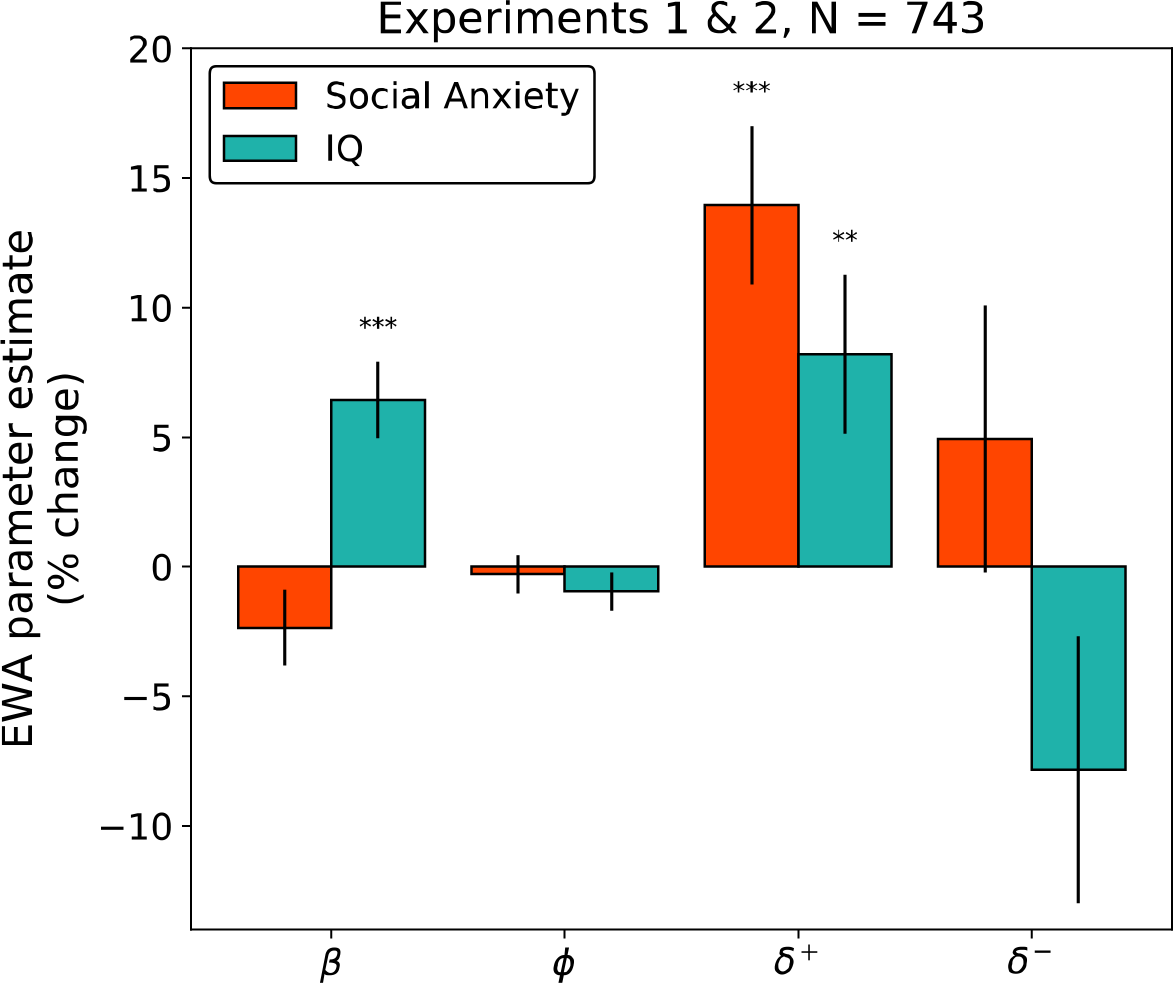
Percent change in valenced EWA parameters as a function of social anxiety (LSAS) and IQ (Abbreviated 9-item Raven’s Matrices) for subjects from Experiments 1 and 2 (N=743). Here *β* represents the inverse temperature, *ϕ* controls the learning rate, and, which dictates the rate of counterfactual updating, is split into *δ*^+^ and *δ*^−^ which control upwards- and downwards- counterfactual learning, respectively. The y-axes indicate the percent change in the parameter for each change of 1 standard deviation (SD) in the predictor. Error bars indicate 1 standard error.

Finally, we examined whether our results were specific to social anxiety by controlling for additional psychopathological symptoms. In general, there are complex patterns of comorbidity among different mental illnesses, and the effects we observed might in principle be subserved by other factors. Notably, rumination is common in other mood disorders, including depression and generalized anxiety. Moreover, a number of studies have linked deficits in goal-directed choice to compulsivity. We used participants’ responses to a larger battery of self-report symptom questionnaires in Experiment 2 to examine how counterfactual reasoning in the patent race task related to symptoms of psychiatric conditions other than social anxiety. We summarized these using the transdiagnostic dimensions identified by Gillan et al. (2016) using factor analysis on the same battery studied here. Using this method, we computed scores for each subject along three dimensions: ‘Anxious-Depression’, ‘Compulsive Behavior and Intrusive Thought’ and ‘Social Withdrawal’ (the last corresponding largely to LSAS; see *SI Figures 5 & 6* for factor loadings). Even controlling for these other factors, upwards counterfactual learning predicted social anxiety (now captured by the ‘Social Withdrawal’ factor; *Figure 5*). The other psychiatric factors did not correlate significantly with any model parameters in this task. Again, the relationship between ‘Social Withdrawal’ and counterfactual updating was significantly greater for upwards counterfactual updating *δ*^+^ vs. downwards counterfactual updating *δ*^−^ (*p* < .03). None of the estimated coefficients for the other terms reached significance (p>.1).

**Figure 5:**
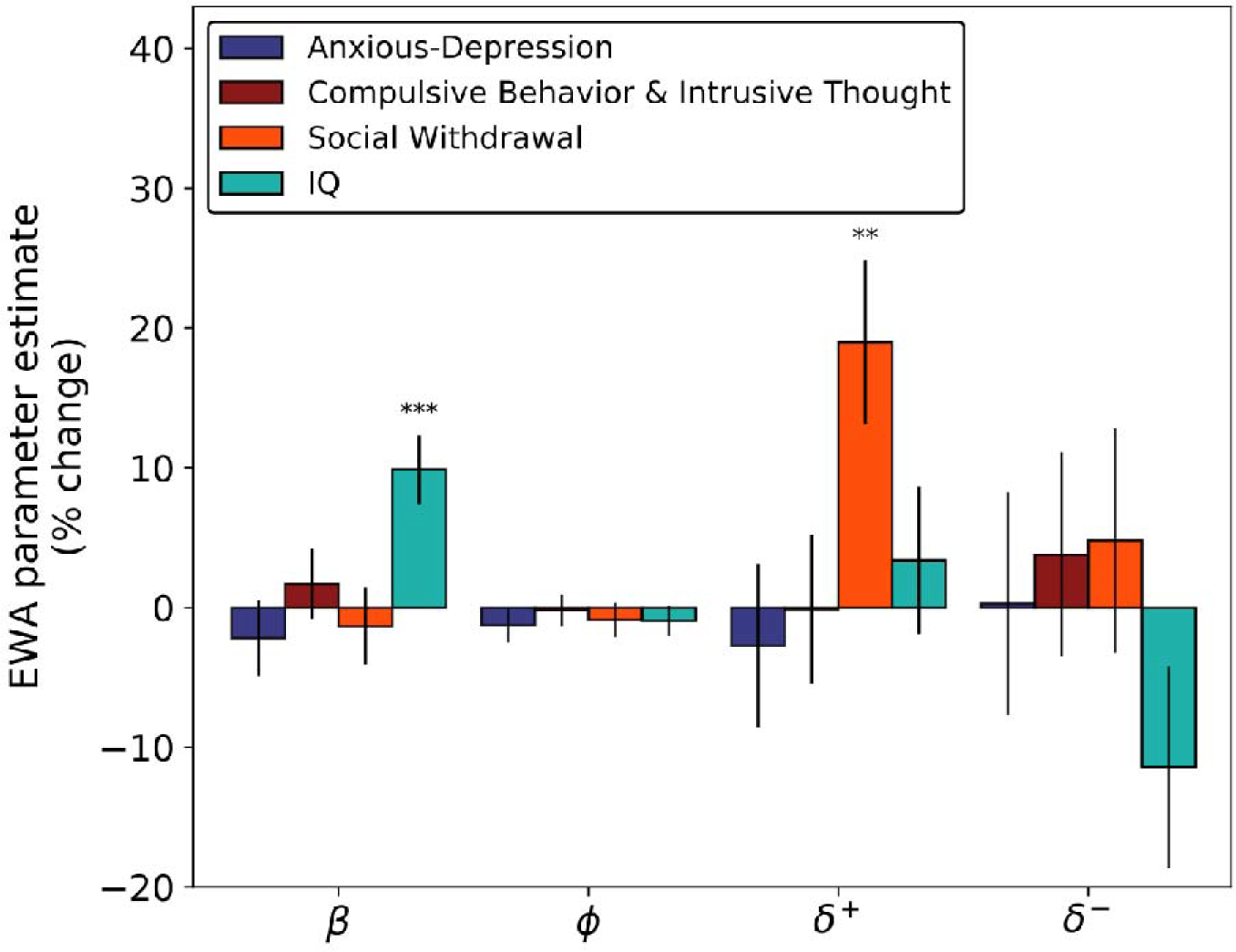
Percent change in EWA parameters as a function of psychiatric symptom dimensions and IQ (Abbreviated 9-item Raven’s Matrices) for Experiment 2 subjects (N = 331). Here *β* represents the inverse temperature, *ϕ* controls the learning rate, and *δ*, which dictates the rate of counterfactual updating, is split into *δ*^+^ and *δ*^−^, which control upwards- and downwards- counterfactual learning, respectively. The y-axes indicate the percent change in the parameter for each change of 1 standard deviation (SD) in the predictor. Error bars indicate 1 standard error.

## Discussion

We investigated the relationship between social anxiety and model-based “belief learning” in two large-scale behavioral experiments. By fitting the parameters of the EWA model to subjects’ choices in a patent race task, we derived indices reflecting the extent to which each participant’s behavior reflected valuation of actions via direct, model-free reinforcement vs. model-based learning of the opponent’s move distribution via counterfactual updating. In line with our hypothesis, self-report social anxiety (LSAS) predicted a significant increase in model usage, as indexed by the EWA parameter *δ*. This effect was driven principally by increased upwards counterfactual reasoning, and robust to the inclusion of additional dimensions of psychiatric symptoms.

These results support the hypothesis that heightened self-reported social anxiety is accompanied by enhanced model-based learning. This finding complements previous reports (albeit using a different, single-player RL task) that symptoms of compulsive disorders are associated with the opposite imbalance: declines in model-based learning (Voon et al., 2015; Gillan et al., 2016). Results of this sort are tantalizing in part because they point the way toward grounding the symptoms of mental illness – here, overthinking and paralysis in social situations – in the dysfunction of well characterized, more basic neuro-computational mechanisms of evaluation and learning. Such a mechanistic understanding of these illnesses – known in medicine as an etiology – is currently wholly lacking, a state of affairs that hampers both diagnosis and treatment, and ultimately contributes to poor patient outcomes. The present study raises the question whether social anxiety is special in this respect, relative to other depressive or anxious disorders. Some evidence that it might be comes from our finding, in Experiment 2, of no similar associations of task behavior with a factor comprising other depression and anxiety symptoms. Of course, we chose a socially framed RL task in order to highlight any effect of social anxiety specifically. The details of the task might well matter: The association of social anxiety with model-basedness was stronger and clearer here than the analogous trend in Gillan et al.’s (2016)’s study of a single-player Markov decision task. Conversely, in the current task we found no evidence that compulsivity is associated with reduced model-based learning, in contrast to Gillan’s setting. The negative finding regarding compulsion is surprising given that OCD in particular has been associated with reduced model-based or goal-directed learning across a range of other tasks, beyond two-step Markov decision tasks (Gillan et al., 2011, 2014); presumably something about how decision making is operationalized in the current task accounts for the difference. (Other than the social framing, differences from the two-step task include that the task is nonsequential and differences in the parameterization of the model, such that the parameter *δ* here isolates the strength of model-based updating, whereas the analogous parameter most strongly affected in two-step tasks, *β*^*MB*^, also incorporates an element of choice consistency, like *β* in the current model.) A related question is why, apart from the social framing, is the effect seen specifically with respect to social anxiety rather than the dimension capturing depressive and other more general anxious symptoms. Social anxiety may indeed be distinct from other mood disorders insofar as overthinking in social contexts is in some sense, goal-directed and task-focused, as opposed to worry and repetitive thought in other disorders which may be more idle and distracting (M. Paulus & M. Stein, personal communication; Watkins, 2008). Social anxiety is also enriched in the sampled population (giving us relatively better power to observe its effects), and simpler and more homogenous as a disorder compared to more complex syndromes like depression. Future work comparing multiple tasks within the same cohort, and manipulating task framing while holding task structure fixed, will be required to fully address these issues.

A broader question is what is the mechanism by which the brain accomplishes “model-based” evaluation – and how is it affected by social anxiety. Our result is consistent with longstanding suggestions that mood disorders might be associated with abnormal or excessive deliberative (model-based) processing, such as uncontrolled forward search for action valuation (Huys et al., 2015; Huys et al., 2012; Solway et al., submitted). For instance, theoretical work on value-based computation in depression has analogized rumination to mental simulation, for computing the value of potential actions given their anticipated consequences, though there has so far been limited evidence connecting these hypothesized computations to actual choices (Huys et al., 2012).

That rumination is, by definition, *selective* – narrowly focused on specific classes of thoughts or events – highlights an important feature of evaluation in general and our data in particular. Much earlier research has viewed “model-based” learning as all-or-nothing, exhaustive recomputation of action values over a tree of future states (Daw et al., 2005) and characterized individual differences in the overall tendency to deploy it (Gillan et al., 2016). But recently, more realistic process-level accounts are emerging that emphasize the selective, strategic contemplation of individual actions or events (Morris & Cushman, 2015; Keramati et al., 2016; Mattar & Daw, 2018). Further, these mechanisms are not limited to online deliberation about decisions presently faced, but also extend to situations like offline planning and anticipatory updating of action values when an outcome is received (Wimmer & Shohamy, 2012; Shohamy & Daw, 2015; Momennejad et al., 2017; Mattar & Daw, 2018).

Suggestively, the EWA model we use is also framed in these terms: Rather than computing action values based on explicit beliefs about the opponent’s move preferences, it achieves the same effect by “counterfactual” updating of moves that could have been chosen, in light of the opponent’s response (Camerer & Ho, 1999). Our finding that social anxiety’s effects on choice were specifically mediated by upwards counterfactual updating speaks directly for this type of computation. Of course, just as *δ* is only one component of learning, *δ*^+^ and *δ*^−^ capture only one dimension of upward and downward comparisons, so our results do not rule out a broader mechanism of valence-dependent computation. Although we do not assess rumination directly (and this is an important direction for future work), this tendency to overweight negative counterfactual prediction errors relative to positive counterfactual prediction errors echoes the general regularities of ruminative content in social anxiety (Kocovski et al., 2005). There is also evidence that people, overall, tend to entertain positive counterfactuals more than negative ones (Kahneman & Miller, 1986; McCloy & Byrne, 2000; Loomes & Sudgen, 1982); It has been argued such a preference is often adaptive for learning (Icard, Cushman, & Knobe, 2018; Mattar & Daw, 2018; Caplin, Dean, & Leahy, 2017). This is also reflected in our data in that baseline *δ*^+^ is greater than *δ*^−^; though we also see that over and above that social anxiety is associated with a further increase in upward counterfactuals, potentially reflecting excessive upward counterfactual rumination that accompanies it. Our initial finding points toward the promise of a better understanding of the microstructure of planning – what events are contemplated, when and why – for understanding and perhaps beginning to address many psychiatric phenomena, including not just compulsivity and rumination but also craving, obsession, and hallucination.

## Acknowledgements

We are grateful to Quentin Huys, Martin Paulus, Murray Stein, and Alec Solway for helpful conversations.

## Supplementary Information

**SI Table 1:** Multivariate linear regressions of social anxiety (LSAS) and IQ (Ravens Matrices Scores) on fitted valenced EWA parameters (N = 743). Here β represents the inverse temperature, *ϕ* controls the learning rate, and *δ*, which dictates the rate of counterfactual updating, is split into *δ*^+^ and *δ*^−^, which control upwards- and downwards- counterfactual learning, respectively.

| DV | IV | Est. (SE) | t-value | Pr>\|t\| |
| --- | --- | --- | --- | --- |
| $\beta$ | Intercept | 0.847 (0.006) | 135.586 | <0.001*** |
|  | Social Anxiety | -0.002 (0.006) | -0.387 | 0.699 |
|  | IQ | -0.008 (0.006) | -1.292 | 0.197 |
| $\phi$ | Intercept | 1.238 (0.018) | 68.259 | <0.001*** |
|  | Social Anxiety | -0.029 (0.018) | -1.608 | 0.108 |
|  | IQ | 0.080 (0.018) | 4.391 | <0.001*** |
| $\delta^+$ | Intercept | 0.286 (0.009) | 32.772 | <0.001*** |
|  | Social Anxiety | 0.040 (0.009) | 4.566 | <0.001*** |
|  | IQ | 0.023 (0.009) | 2.684 | 0.007** |
| $\delta^-$ | Intercept | 0.136 (0.007) | 19.419 | <0.001*** |
|  | Social Anxiety | 0.007 (0.007) | 0.955 | 0.340 |
|  | IQ | -0.011 (0.007) | -1.520 | 0.1288 |

**SI Table 2:** Multivariate linear regressions of psychiatric symptom dimensions (factors) and IQ (Ravens Matrices Scores) on fit valenced EWA parameters. Factors were ‘Anxious-Depression’, ‘Compulsive Behavior and Intrusive Thought’ and ‘Social Withdrawal’ (Experiment 2, N = 331). Here *β* represents the inverse temperature, *ϕ* controls the learning rate, and *δ*, which dictates the rate of counterfactual updating, is split into *δ*^+^ and *δ*^−^, which control upwards- and downwards- counterfactual learning, respectively.

| DV | IV | Est. (SE) | t-value | Pr>\|t\| |
| --- | --- | --- | --- | --- |
| $\beta$ | Intercept | 1.205 (0.029) | 40.875 | <0.001*** |
|  | Anxious Depression | -0.026 (0.033) | -0.805 | 0.422 |
|  | Compulsive Behavior & Intrusive Thought | 0.020 (0.030) | 0.674 | 0.501 |
|  | Social Withdrawal | -0.016 (0.033) | -0.490 | 0.625 |
|  | IQ | 0.106 (0.026) | 4.007 | <0.001*** |
| $\phi$ | Intercept | 0.847 (0.009) | 90.391 | <0.001*** |
|  | Anxious Depression | -0.011 (0.010) | -1.048 | 0.296 |
|  | Compulsive Behavior & Intrusive Thought | -0.002 (0.009) | -0.178 | 0.859 |
|  | Social Withdrawal | -0.008 (0.010) | -0.732 | 0.465 |
|  | IQ | -0.007 (0.008) | -0.865 | 0.388 |
| $\delta^+$ | Intercept | 0.256 (0.013) | 19.009 | <0.001*** |
|  | Anxious Depression | -0.007 (0.015) | -0.469 | 0.639 |
|  | Compulsive Behavior & Intrusive Thought | 0.000 (0.014) | -0.023 | 0.982 |
|  | Social Withdrawal | 0.049 (0.015) | 3.232 | 0.001** |
|  | IQ | 0.008 (0.012) | 0.640 | 0.523 |
| $\delta^-$ | Intercept | 0.155 (0.011) | 13.719 | <0.001*** |
|  | Anxious Depression | 0.001 (0.013) | 0.035 | 0.972 |
|  | Compulsive Behavior & Intrusive Thought | 0.006 (0.011) | 0.519 | 0.604 |
|  | Social Withdrawal | 0.008 (0.013) | 0.596 | 0.552 |
|  | IQ | -0.016 (0.010) | -1.588 | 0.113 |

**SI Table 3:**
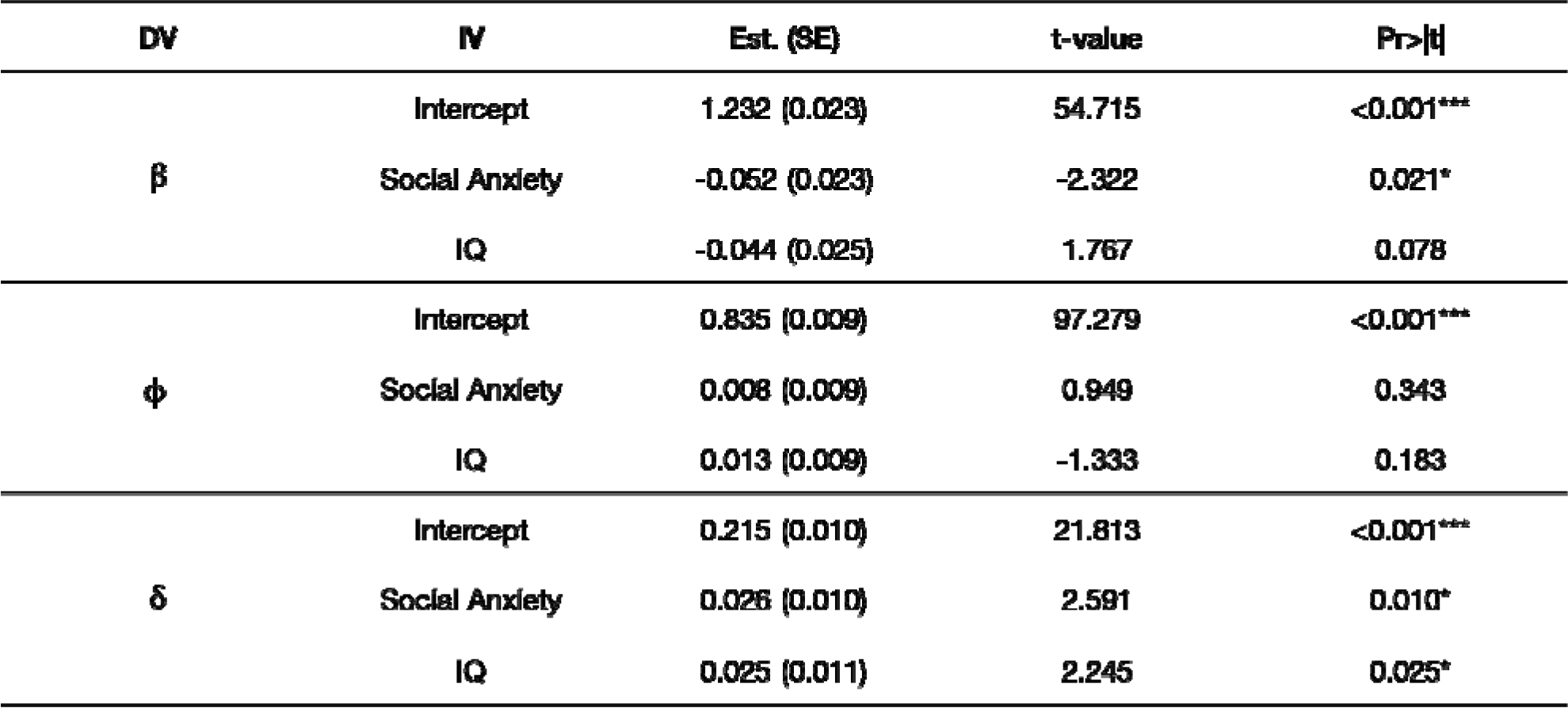
Multivariate linear regressions of social anxiety (LSAS) and IQ (Ravens Matrices Scores) on fitted EWA parameters (Experiment 1, N = 412). Here *β* represents the inverse temperature, *ϕ* controls the learning rate, and *δ* dictates the relative rate of counterfactual updating.

| DV | IV | Est. (SE) | t-value | Pr>\|t\| |
| --- | --- | --- | --- | --- |
| $\beta$ | Intercept | 1.232 (0.023) | 54.715 | <0.001*** |
|  | Social Anxiety | -0.052 (0.023) | -2.322 | 0.021* |
|  | IQ | -0.044 (0.025) | 1.767 | 0.078 |
| $\phi$ | Intercept | 0.835 (0.009) | 97.279 | <0.001*** |
|  | Social Anxiety | 0.008 (0.009) | 0.949 | 0.343 |
|  | IQ | 0.013 (0.009) | -1.333 | 0.183 |
| $\delta$ | Intercept | 0.215 (0.010) | 21.813 | <0.001*** |
|  | Social Anxiety | 0.026 (0.010) | 2.591 | 0.010* |
|  | IQ | 0.025 (0.011) | 2.245 | 0.025* |

**SI Table 4:** Multivariate linear regressions of social anxiety (LSAS) and IQ (Ravens Matrices Scores) on fitted EWA parameters (Experiment 2, N = 331). Here *β* represents the inverse temperature, *δ* controls the learning rate, and *δ* dictates the relative rate of counterfactual updating.

| DV | IV | Est. (SE) | t-value | Pr>\|t\| |
| --- | --- | --- | --- | --- |
| $\beta$ | Intercept | 1.135 (0.035) | 32.737 | <0.001*** |
|  | Social Anxiety | -0.001 (0.034) | -0.023 | 0.981 |
|  | IQ | 0.107 (0.031) | 3.521 | <0.001*** |
| $\phi$ | Intercept | 0.835 (0.010) | 86.498 | <0.001*** |
|  | Social Anxiety | -0.016 (0.010) | -1.677 | 0.095 |
|  | IQ | -0.009 (0.008) | -1.053 | 0.293 |
| $\delta$ | Intercept | 0.214 (0.014) | 15.811 | <0.001*** |
|  | Social Anxiety | 0.034 (0.013) | 2.507 | 0.013* |
|  | IQ | 0.002 (0.012) | 0.169 | 0.866 |

**SI Table 5:**
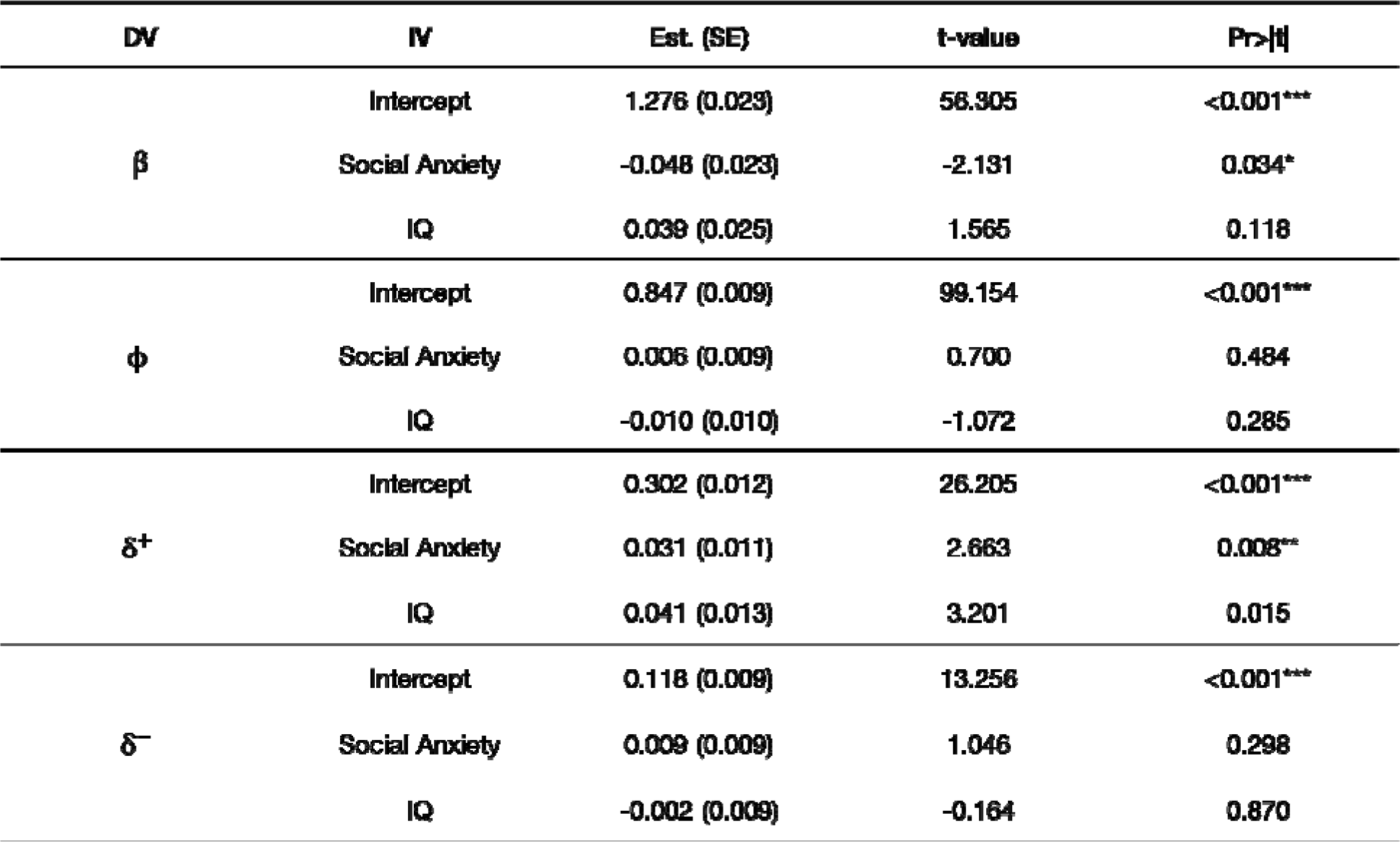
Multivariate linear regressions of social anxiety (LSAS) and IQ (Ravens Matrices Scores) on fitted valenced EWA parameters (Experiment 1, N = 412). Here *β* represents the inverse temperature, *ϕ* controls the learning rate, and *δ* dictates the relative rate of counterfactual updating.

**SI Table 6:**
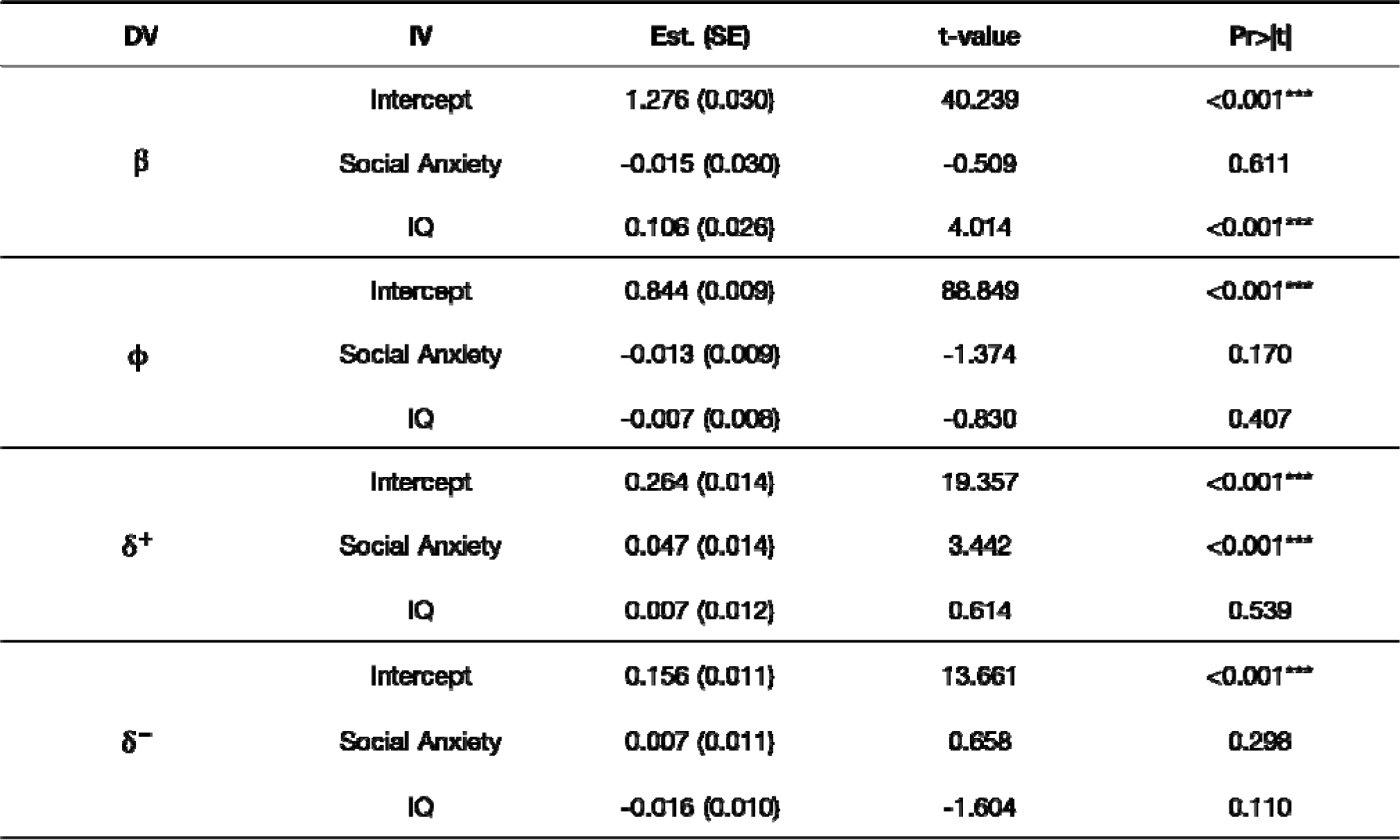
Multivariate linear regressions of social anxiety (LSAS) and IQ (Ravens Matrices Scores) on fitted valenced EWA parameters (Experiment 2, N = 331).

**SI Table 7:** Parameter fit values for the standard (top) and valenced (bottom) versions of the EWA model estimated separately for each experiment. Here *β* represents the inverse temperature, *ϕ* controls the learning rate, and *δ*, which dictates the rate of counterfactual updating, is split into *δ*^+^ and *δ*^−^, which control upwards- and downwards- counterfactual learning, respectively. The columns labeled and refer respectively to the mean and standard deviations (in parentheses) of the subject-level fit values for each experiment.

| Model | Parameter | Experiment 1 |  | Experiment 2 |  |
| --- | --- | --- | --- | --- | --- |
| | | $\mu$ | ($\sigma$) | $\mu$ | ($\sigma$) |
| EWA | $\beta$ | 1.2 | (0.46) | 1.1 | (0.63) |
| | $\phi$ | 0.84 | (0.17) | 0.84 | (0.17) |
| | $\delta$ | 0.22 | (0.20) | 0.21 | (0.24) |
| | $V^1(1)$ | 0.013 | (0.0093) | -0.053 | (0.019) |
| | $V^2(1)$ | -0.34 | (0.019) | -0.26 | (0.0078) |
| | $V^3(1)$ | -0.11 | (0.029) | -0.17 | (0.0078) |
| | $V^4(1)$ | 0.078 | (0.015) | 0.09 | (0.043) |
| | $V^5(1)$ | 0.13 | (0.012) | 0.15 | (0.014) |
| Valenced EWA | $\beta$ | 1.3 | (0.46) | 1.2 | (0.55) |
| | $\phi$ | 0.85 | (0.17) | 0.85 | (0.17) |
| | $\delta^+$ | 0.31 | (0.24) | 0.26 | (0.25) |
| | $\delta^-$ | 0.12 | (0.18) | 0.16 | (0.21) |
| | $V^1(1)$ | 0.015 | (0.0093) | -0.044 | (0.014) |
| | $V^2(1)$ | -0.34 | (0.020) | -0.27 | (0.013) |
| | $V^3(1)$ | -0.11 | (0.024) | -0.17 | (0.012) |
| | $V^4(1)$ | 0.083 | (0.014) | 0.11 | (0.017) |
| | $V^5(1)$ | 0.13 | (0.011) | 0.16 | (0.014) |

**SI Table 8:** Comparison of Nash equilibrium predictions and empirical distributions from Rapoport & Amaldoss (2000), Zhu et al., (2012), and Experiments 1 & 2.

**Empirical Distributions of Investment Strategies**
|  | Investment |  |  |  |  |
| --- | --- | --- | --- | --- | --- |
| | \$0 | \$1 | \$2 | \$3 | \$4 |
| Nash Equilibrium | 60% | 0% | 20% | 0% | 20% |
| Rapoport & Amaldoss (2000) | 55% | 3% | 6% | 14% | 22% |
| Zhu et al., (2012) | 49% | 3% | 10% | 10% | 28% |
| Experiments 1 & 2 | 41% | 6% | 9% | 13% | 31% |

**SI Figure 3:**
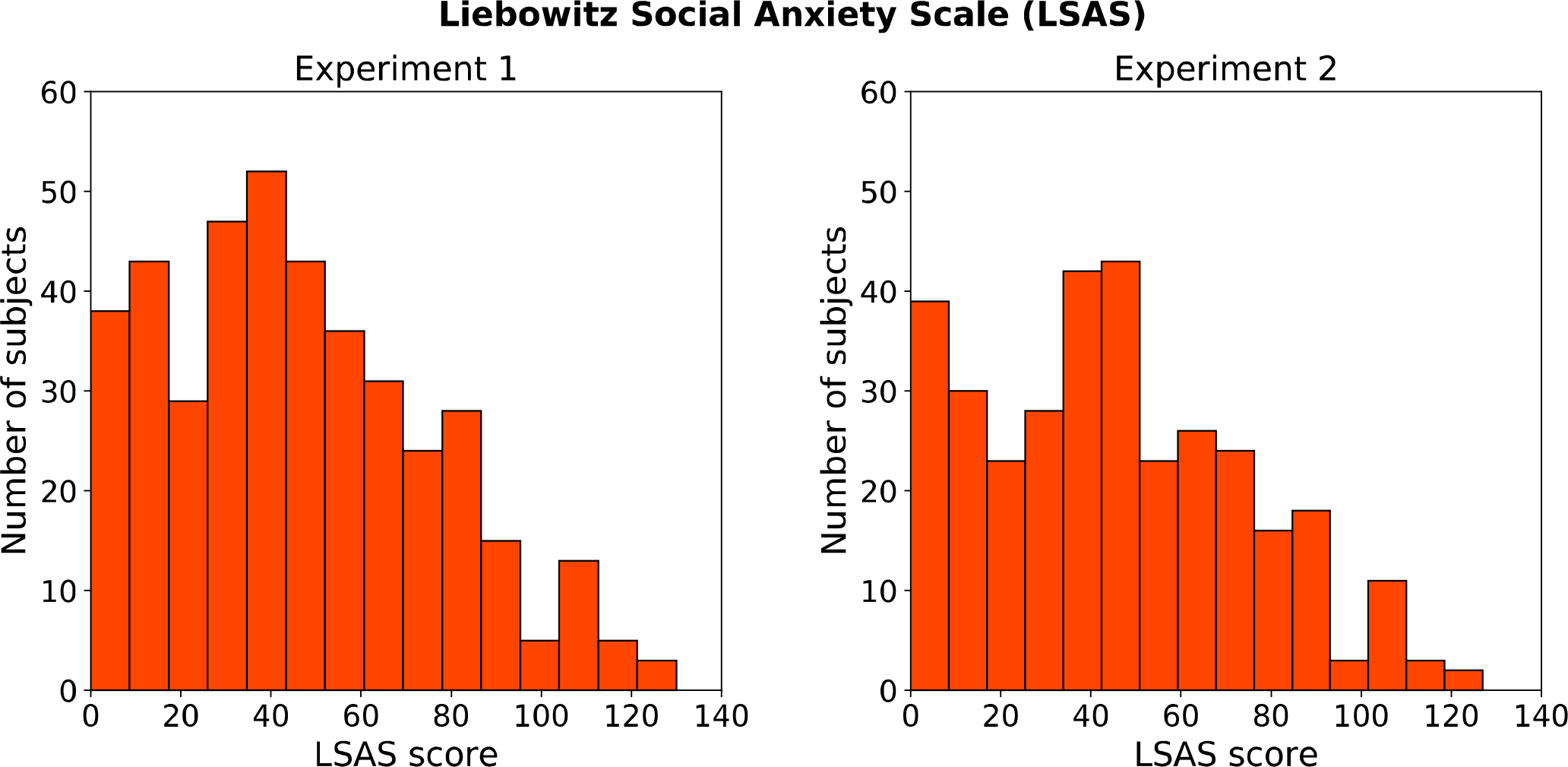
Distributions of subjects’ scores on the Liebowitz Social Anxiety Scale (LSAS) for Experiment 1 (left) and Experiment 2 (right).

**SI Figure 4:**
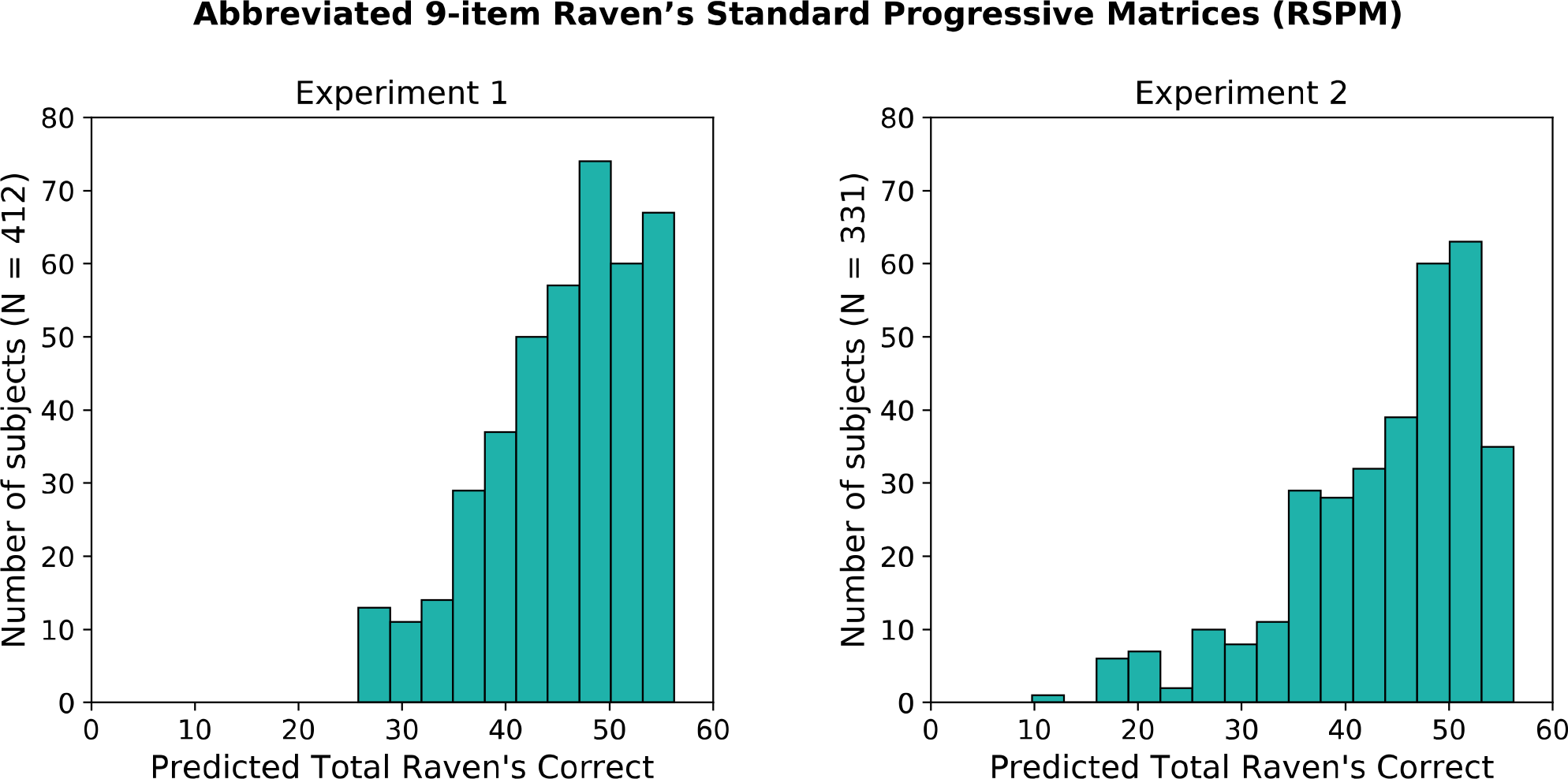
Projected IQ scores for Experiments 1 (left) and Experiment 2 (right) based on subjects’ responses to the 9- item abbreviated version of the Raven’s Standard Progressive Matrices (RSPM).

**SI Figure 5:**
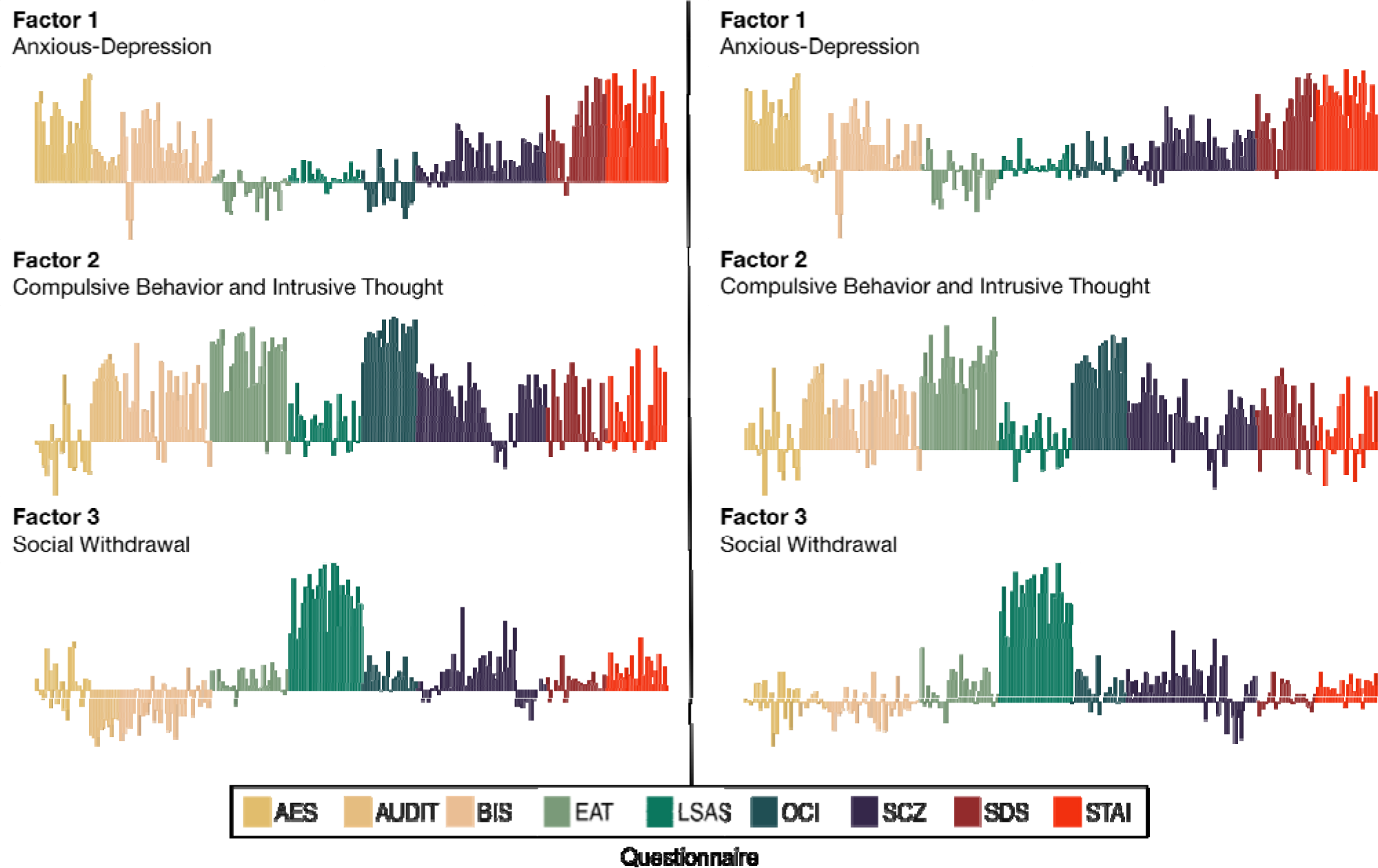
Item loadings for (trans-diagnostic) psychiatric factors derived from data in Gillan et al., (2016), (N= 1413) (left panel) and Experiment 2, (N = 331) (right panel). Factor analysis on the correlation matrix of 209 questionnaire items suggested that 3-factor solution best explained these data. Factors were ‘Anxious-Depression’, ‘Compulsive Behavior and Intrusive Thought’ and ‘Social Withdrawal’. Item loadings for each factor are indicated by the height of the vertical bars for each respective item (bars extending downwards indicate negative loadings). For convenience, item bars are grouped by questionnaire along each factor’s horizontal axis. the color-code (bottom) specifies the questionnaire from which each item was drawn: AES (Apathy Evaluation Scale), AUDIT (Alcohol Use Disorders Identification Test), BIS (Barratt Impulsiveness Scale 11), EAT (Eating Attitudes Test), LSAS (Liebowitz Social Anxiety Scale), OCI (Obsessive-Compulsive Inventory-Revised [OCI-R]), SCZ ((Short Scales for Measuring Schizotypy), SDS (Zung Self-Rating Depression Scale), STAI (State Trait Anxiety Inventory). As implied by the shared resemblance of the left and right panels, the factor loadings derived from the two datasets were highly correlated: Factor 1: R = .94, p <1e-96; Factor 2: R = .91, p <1e-79; Factor 3: R = .91, p <1e-80).

**SI Figure 6:**
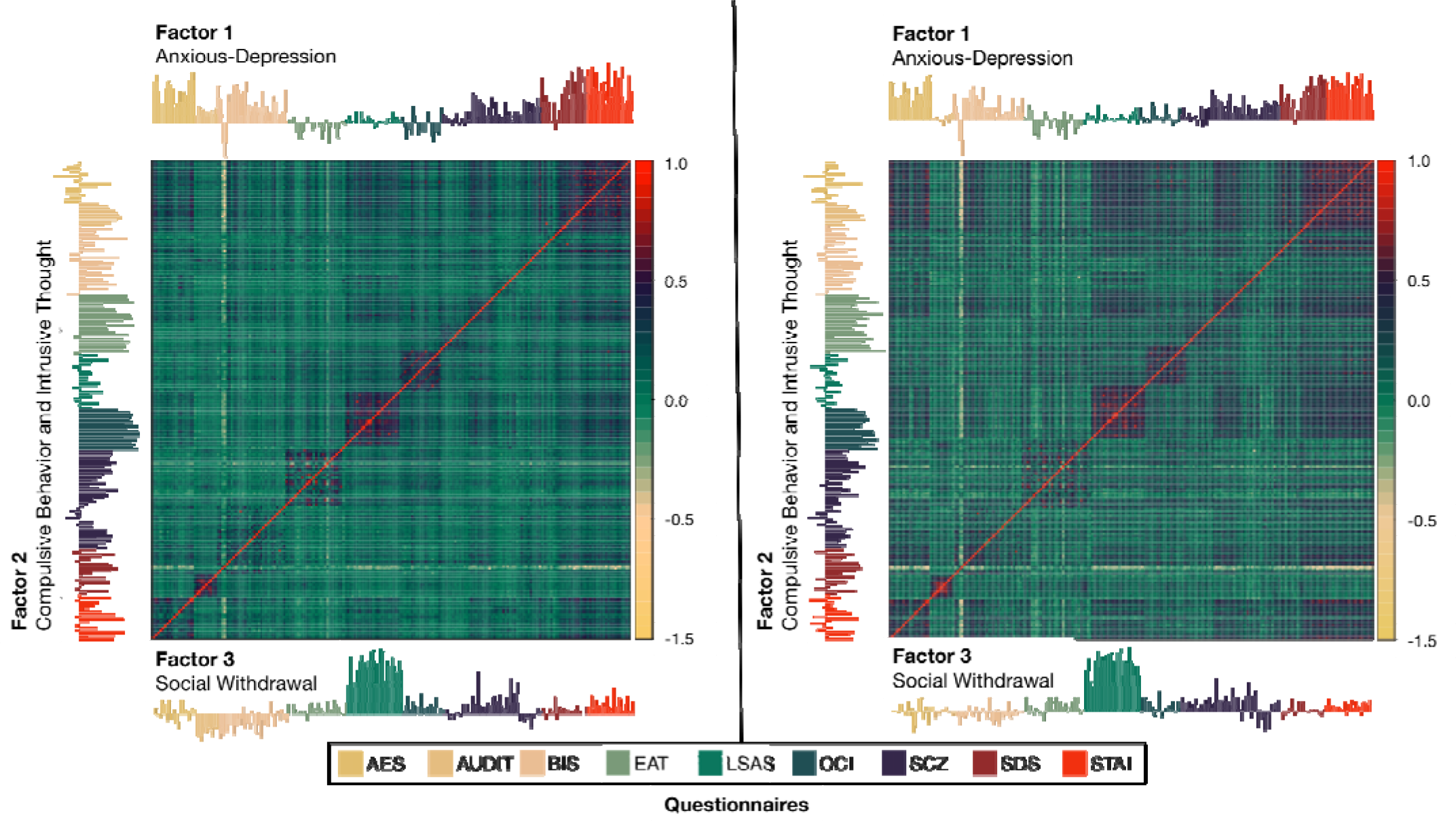
Correlation matrices of 209 questionnaire items based on data in Gillan et al., (2016), (N= 1413) (left) and Experiment 2, (N = 331) (right). Factor analysis indicated that these data were best explained by 3- factors: ‘Anxious-Depression’, ‘Compulsive Behavior and Intrusive Thought’ and ‘Social Withdrawal’. Items are grouped by questionnaire (indicated by the color-code) and the individual item loadings for each factor for are presented on the top, left and bottom sides of the correlation matrix.

**SI Figure 7:**
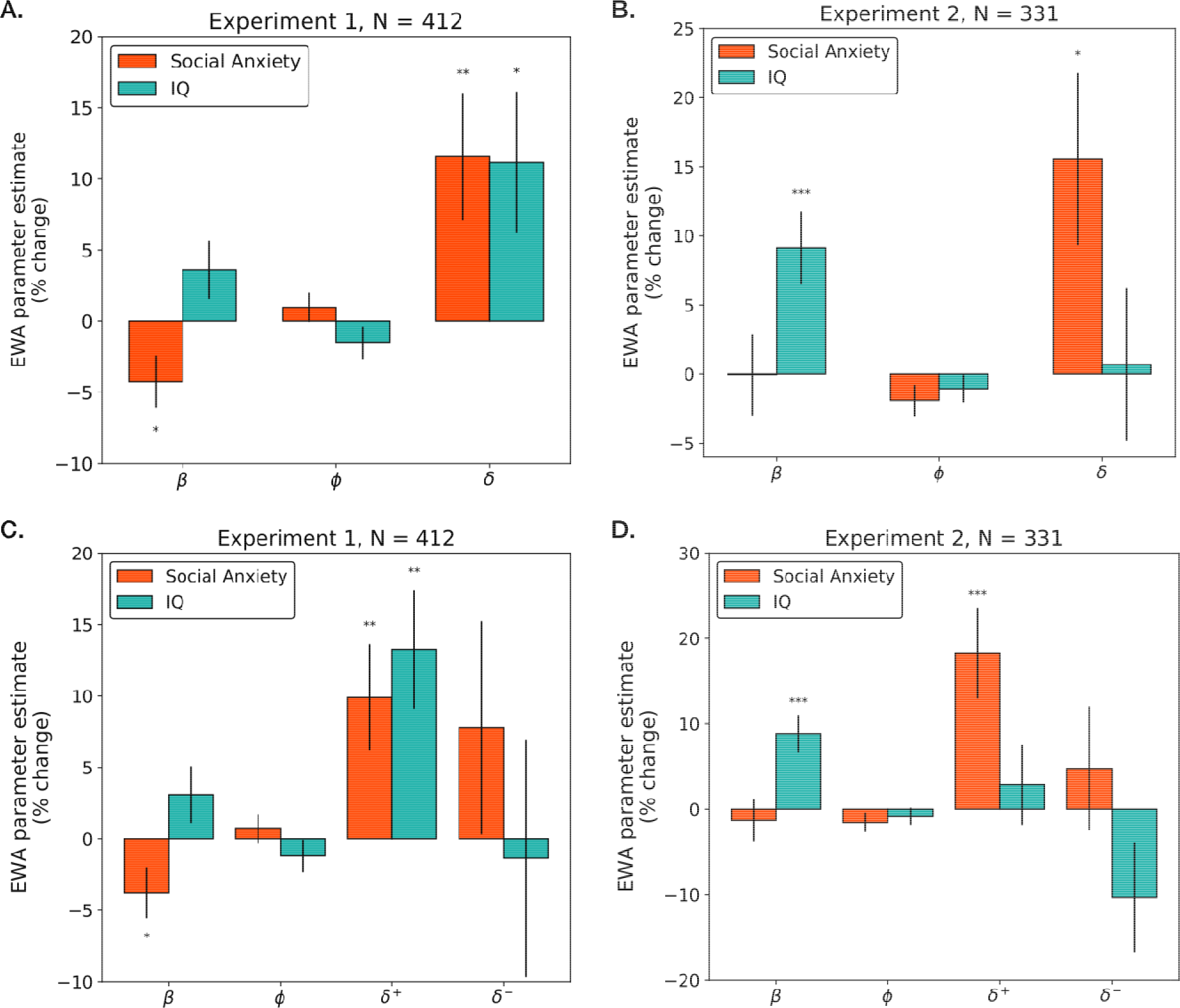
Percent change in EWA parameters as a function of social anxiety (LSAS) and IQ (Abbreviated 9-item Raven’s Matrices) for subjects from Experiment 1 (A, C; N=412) and Experiment 2 (B, D; N = 331). The y-axes indicate the % change in the dependent variable (i.e., *β*, *ρ*) for each change of 1 standard deviation (SD) in the predictor (i.e., LSAS or IQ) and error bars indicate standard error. The upper and lower panels depict parameters estimated according to the standard EWA model (A. and B.) and the valenced EWA model (C. and D.) respectively.

